# Neural structure mapping in human probabilistic reward learning

**DOI:** 10.1101/366757

**Authors:** F. Luyckx, H. Nili, B. Spitzer, C. Summerfield

## Abstract

Humans can learn abstract concepts that describe invariances over relational patterns in data. One such concept, known as magnitude, allows stimuli to be compactly represented by a single dimension (i.e. on a mental line), for example according to their cardinality, size or value. Here, we measured representations of magnitude in humans by recording neural signals whilst they viewed symbolic numbers. During a subsequent reward-guided learning task, the neural patterns elicited by novel complex visual images reflected their pay-out probability in a way that suggested they were encoded onto the same mental number line. Our findings suggest that in humans, learning about values is accompanied by structural alignment of value representations with neural codes for the concept of magnitude.

---

The ability to learn rapidly from limited data is a key ingredient of human intelligence. For example, on moving to a new city, you will rapidly discover which restaurants offer good food and which neighbors provide enjoyable company. Current models of learning propose that appetitive actions towards novel stimuli are learned *tabula rasa* via reinforcement (*1*), and these models explain the amplitude of neural signals in diverse brain regions during reward-guided choices in humans and other animals (*2–4*). However, reinforcement learning models learn only gradually, and even when coupled with powerful function approximation methods, exhibit limited generalization beyond their training domain (*5*), leading to the suggestion they are ill-equipped to fully describe human learning (*6*). By contrast, cognitive scientists have ascribed human intelligence to formation of abstract knowledge representations (or “concepts”) that delimit the structural forms that new data is likely to take (*7–9*). Indeed, real-world data can often be described by simple relational structures, such as a tree, a grid or a ring (*7*), and humans may infer relational structure through probabilistic computation (*10*) and understand new domains by their alignment with existing relational structures (*11*). However, these models are often criticized for failing to specify how concepts might be plausibly encoded or computed in neural circuits (*12*). A pressing concern, thus, is to provide a mechanistic account of how relational knowledge is encoded and generalized in the human brain (*13*).

The current project was inspired by recent observations that the representational geometry of human neural signals evoked by symbolic numbers respects their relative cardinality (*14, 15*). In scalp M/EEG signals, neural patterns evoked by Arabic digits vary continuously with numerical distance, such that multivariate signals for “3” are more similar to those for “4” than “5”. Number is a symbolic system that expresses magnitude in abstract form (*16–18*) and so we reasoned that continuously-varying neural signals evoked by numbers might be indexing a conceptual basis set that supports one-dimensional encoding of novel stimuli. In the domain of reward-guided learning, a compact description of the stimulus space projects data into a single dimension that runs from “bad” to “good”. Here, thus, we asked humans to learn the reward probabilities associated with novel, high-dimensional visual images, and measured whether the stimuli come to elicit neural patterns that map onto one-dimensional neural codes for numerical magnitude.

Whilst undergoing scalp EEG recordings, human participants (n = 46) completed two tasks: a numerical decision task and a probabilistic reward-guided learning task. In the numerical task participants viewed rapid streams of ten Arabic digits (1 to 6) and reported whether numbers in orange or blue font had the higher (n = 22) or lower (n = 24) average (Fig. 1A). Using representational similarity analysis (RSA) (*19*) we replicated the previous finding (*14, 15*) that patterns of neural activity across the scalp from ~100ms onwards were increasingly dissimilar for digits with more divergent magnitude, i.e. codes for “3” and “5” were more dissimilar than those for “3” and “4” (Fig. 1A, green line). This occurred irrespective of task framing (report higher vs. lower average) and category (orange vs. blue numbers), suggesting that neural signals encoded an abstract representation of magnitude and not solely a decision-related quantity such as choice certainty (*14*). The reward-learning task was based on the multi-armed bandit paradigm that has been used ubiquitously to study value-guided decision-making (*20*). Participants learned the reward probabilities associated with six unique novel images (colored donkeys; Fig. 1A), which paid out a fixed reward with a stationary probability (range 0.05-0.95). These probability values were never signaled to the participant but instead acquired by trial and error learning in an initial learning phase. In the test phase, we asked participants to decide between two successive donkeys to obtain a reward, and estimated trial-wise subjective probability estimates for each bandit by fitting a delta-rule model to choices (*1*). Throughout these phases, the bandits were never associated with numbers in any way.

**Fig. 1.**
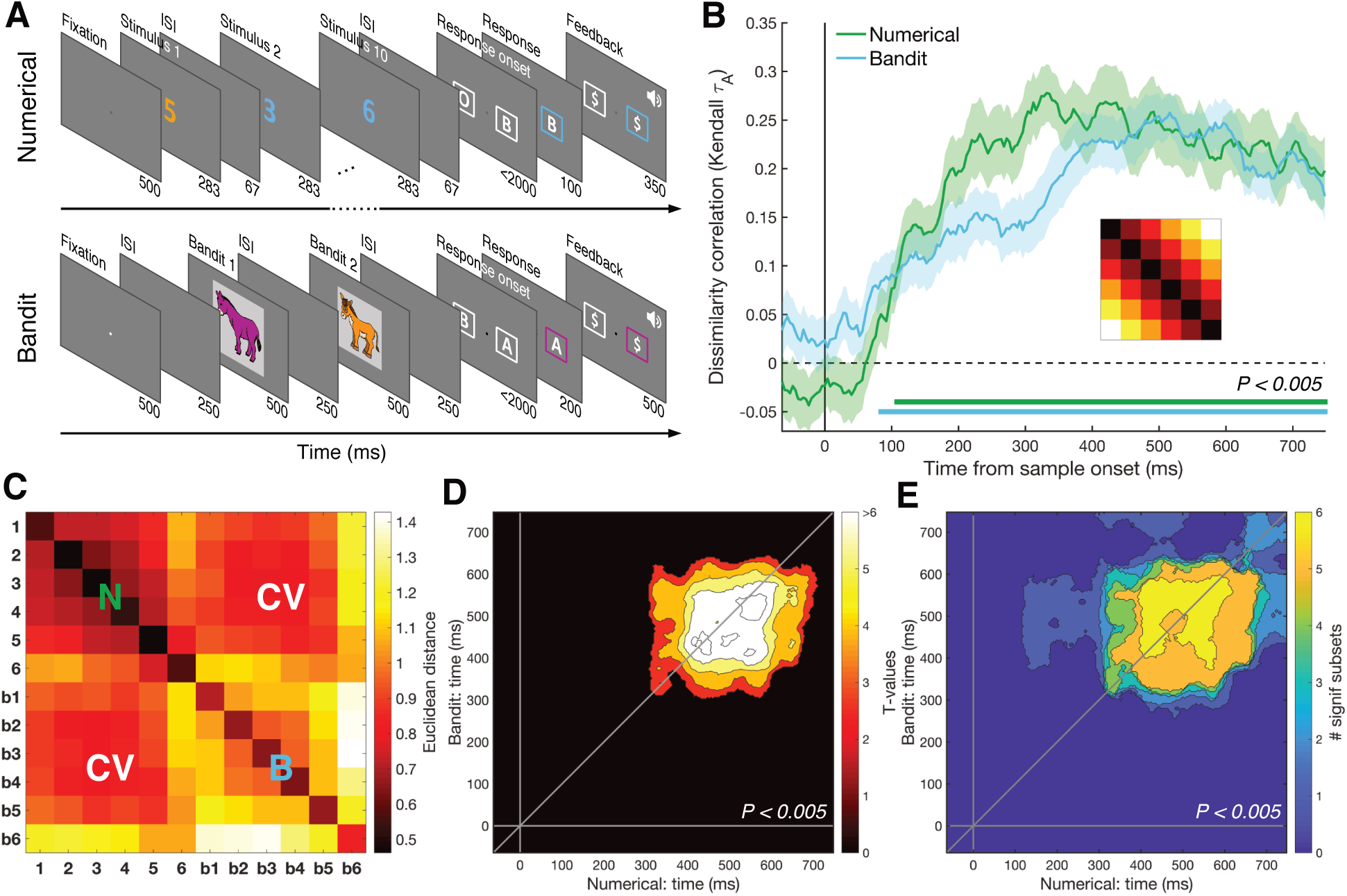
Task design and RSA results. **(A)** Humans performed two tasks during a single EEG recording session. In the numerical decision task, participants viewed a stream of ten digits between 1 and 6, deciding whether the blue or orange numbers had the highest/lowest average. In the bandit task, participants learned about the reward probabilities of six images (bandits) and were asked to choose between two successive bandits to obtain a fixed reward. Numbers below each frame show ISI in ms. **(B)** RSA revealed a numerical and value distance effect from ~100 ms after stimulus onset. Inset shows magnitude model RDM. Shaded area represents SEM. **(C)** Averaged full RDM from 350-600 ms for numbers (1-6) and bandits (b1-b6). Upper left and lower right quadrants show representational similarity for numbers (N) and bandits (B) respectively, i.e. within-task RDM; lower left/upper right quadrants show cross-validated similarity between numbers and bandits (CV), i.e. between-task RDM. **(D)** Cross-temporal cross-validated RDM revealed a stable magnitude representation that was shared between the two tasks. **(E)** This effect was robust to the elimination of any single number/bandit pair. Correction for multiple comparisons was performed with cluster-based permutation tests, P_cluster_ < 0.005.

Next, we used RSA to examine the neural patterns evoked by bandits, ordered by their subjective reward probability. We found that multivariate EEG signals varied with subjective bandit ranks, with bandits that paid out with nearby probabilities eliciting more similar neural patterns (from ~100ms onwards; Fig. 1B, blue line). Our key question was whether there was a shared neural code for numerical magnitude and reward probability. We found that this was the case. For example, EEG signals elicited by digit “6” were more similar to those evoked by the most valuable bandit, and number 1 predicted the bandit least likely to pay out, with a similar convergence for intermediate numbers and bandits (Fig. 1C). Cross-validation of neural signals elicited by all numbers (1 to 6) and bandits (inverse ranks 1-6) was stable and reliable from 300-650 ms post-stimulus, as demonstrated by cross-temporal RSA (Fig. 1D) (*21*). This cross-validated pattern remained robust to the removal of any one of the six number/bandit pairs (Fig. 1E) or exclusion of the diagonal elements (Fig. S1A) and was more consistent within than between participants (Fig. S1B).

To explore how these representations might be learned and generalized at the mechanistic level, we turned to a simple computational tool, a feedforward neural network (*22*). Unlike handcrafted probabilistic models, neural networks are not constrained to make inferences over structure, but structured representations may emerge naturally in the weights during training (*12, 23*). Here, as a proof of concept, we confronted the network with two different stimulus sets in turn that (like our numbers and bandits) shared the same similarity structure. We then asked if the shared structure facilitates retraining on the second set after learning the first (Fig. 2A). The network was first trained on inputs *x*_1_ arriving at input units *X_A_*, and after convergence, retrained on inputs *x_b_* fed into units *X_B_* (where *X_A_* and *X_B_* are separate input modules that project to a common hidden layer H). Inputs *x_b_* were 6 random vectors constructed to have the same continuously-varying similarity structure as the bandits, whereas inputs *x_a_* consisted of either a different set of 6 random vectors with the same second-order structure, or a shuffled control. Relearning on *x_b_* proceeded faster when inputs shared a common structure with *x_a_* (Fig. 2B-D) and RSA conducted after retraining revealed reliable cross-validated patterns of activity in the hidden units for this condition alone (Fig. 2E), mirroring the result from the human neural data.

**Fig. 2.**
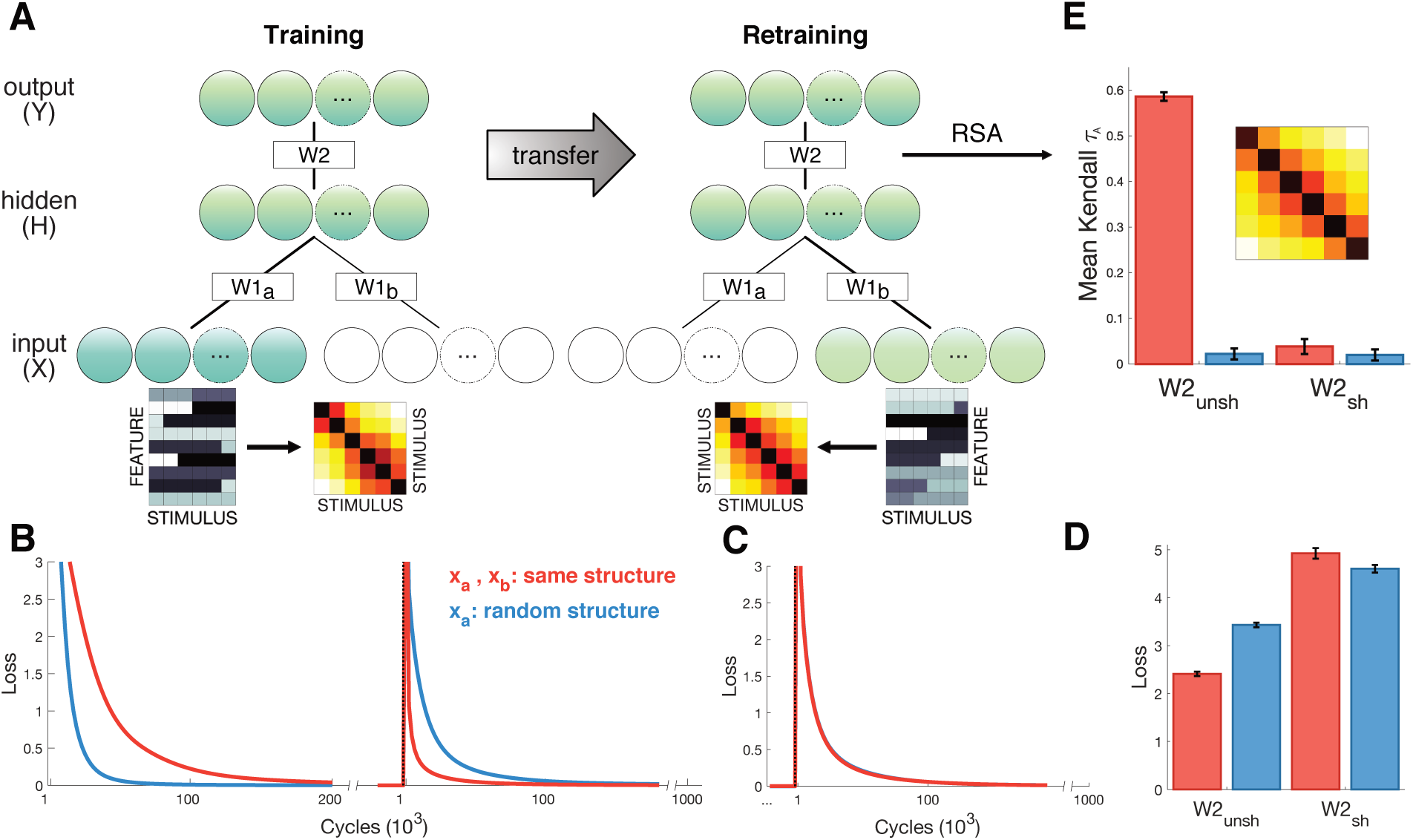
Neural network simulations. **(A)** Schematic depiction of network structure and training. The network was first trained to classify inputs x_a_ fed into units X_A_ (lower blue circles). Inputs x_a_ consisted of 6 stimuli that either exhibited gradual increasing dissimilarity or were shuffled as a control (stimulus RDMs shown next to examples of x_a_ and x_b_). After convergence, the model was trained on new input x_b_ that were fed into a separate input stream X_B_ (lower green circles). Inputs x_b_ were different to x_a_ but exhibited the same similarity structure. **(B)** Loss plotted over the course of training (left panel) and retraining (right panel) for the test (red) and shuffled control (blue) conditions. Learning was faster during training for control stimuli, but retraining was faster when x_a_ and x_b_ exhibited shared similarity structure. **(C)** Loss for control simulations where W2 were shuffled between training and retraining, suggesting successful transfer depends on structure encoded in W2. **(D)** Mean loss for first 1000 cycles after retraining. **(E)** Cross-validation RSA on hidden unit activation for all stimuli in x_a_ and x_b_ after retraining. Hidden unit activations exhibit shared similarity structure only when W2 remains unshuffled and x_a_ and x_b_ share structure.

Our participants were all numerate adults with an intact sense of magnitude, and thus an equivalent “shuffled” control was unavailable for the human data. Nevertheless, building on past work that has described choice biases in numerical cognition and economic decisions (*24*), we asked whether individual differences in the mental number representation explained variance in the neural codes elicited by the bandits. We previously observed that participants overweighted larger magnitudes during the numerical decision task (*14*), e.g. numbers “5” and “6” had disproportionate impact on averaging judgments, and this finding was replicated in the current data (Fig. 3A). Human choices were best fit by a power-law model in which participants averaged and compared distorted numerical values *x^k^* with *k* = 2.04 ± 1.11 (*k* > 1: t(45) = 12.47, P < 0.001). Turning to the neural data, we generated candidate representational dissimilarity matrices (RDMs) under the assumption that distance in neural space can like-wise be non-linear and best described by a distortion of the form *x^k^*. We found that in both tasks the best fitting RDM was parameterized by *k* > 1 [numerical: *k* = 1.73 ± 0.82, t(45) = 14.34, P < 0.001; bandit: *k* = 1.72 ± 0.85, t(45) = 13.65, P < 0.001] (Fig. 3B). Moreover, estimated distortions for behavior and EEG in the numerical task were highly correlated, suggesting that they were driven by shared distortions in neural magnitude coding. To assess whether a distortion in the number line was also associated with choice behavior in the bandit task, we predicted neural patterns in the bandit task from the choice distortions in the numerical task (after partialling out choice distortions estimated from the bandit task itself). Individual differences in warping of the number representation continued to explain variance in the neural bandit representation (Fig. 3C). This implies that humans used their intrinsic sense of magnitude when forming neural representations for the bandits in one-dimensional probability space.

**Fig. 3.**
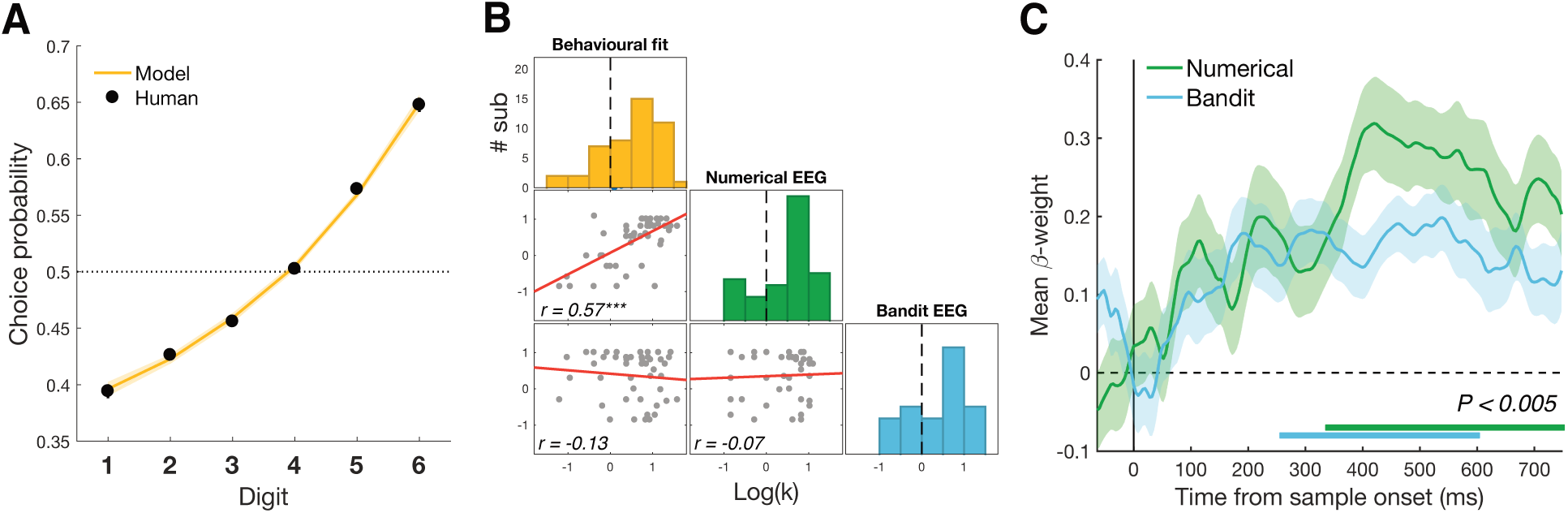
Psychometric and neurometric distortions. **(A)** Choice probabilities in the numerical task were modelled with a psychometric model that transformed input values into a decision value with exponent k. Participant’s responses were best modelled by a distortion parameter k > 1. **(B)** Distribution of estimated k for behavior in the numerical task and neural RDMs for both tasks. All measures were best described by k > 1. Estimated distortions for behavior and EEG in the numerical task were highly correlated. **(C)** Choice distortions from the numerical task explained variance in the neural RDMs of the bandit task after partialling out choice behavior in the bandit task using competitive GLM (P_cluster_ < 0.005).

Recent work has suggested that during categorization, posterior parietal neurons in the macaque monkey are strikingly low-dimensional, as if the parietal cortex were engaging in a gain control process that projected stimulus features or timings on a single axis (*25–27*). Indeed, focusing on centro-parietal electrodes, we observed a univariate positivity that varied with the magnitude of both numbers and reward probabilities (Fig. 4A-B). This signal resembles a previously described EEG signal, known as the CPP, that has been found to scale with the choice certainty in perceptual (*28*) and economic tasks (*29*). However, we note that in our numerical task the CPP followed an approximately ascending pattern from lower to higher numbers regardless of task framing or color category, and that the cross-validation effect persisted even after the CPP had been regressed out of the data (Fig. S2). This suggests that (a) the CPP in our task may represent a notion of magnitude, not a certainty signal; and (b) that this signal is not the sole driver of our multivariate findings. Nevertheless, to understand the dimensionality of the number and bandit representations (and the subspace in which they aligned), we used two dimensionality reduction techniques, singular value decomposition (SVD) and multidimensional scaling (MDS). First, using SVD, we systematically removed dimensions from the EEG data and recomputed our number-bandit cross-validation scores. We found that probabilistic reward learning was supported by a low-dimensional neural magnitude code, with reliable effects persisting when all but 2 eigenvectors were removed from the data but attenuated when only a single dimension was retained in the EEG data. Statistical comparison suggested that within the 350-600 ms period for which significance was observed, the 2D and full (high-dimensional) solutions led to equivalent cross-validation but the reduction to 1D significantly reduced the effect (Fig. 4C). Secondly, we used MDS to visualize the first dimensions of the concatenated number/bandit data. This disclosed an axis pertaining to magnitude (Fig. 4D) and another for certainty along which, especially for the bandits, the large (or best) and small (or worst) items diverged from the others. In other words, the numbers and bandits align principally along a single magnitude axis but with an additional contribution from a second factor encoding choice certainty.

**Fig. 4.**
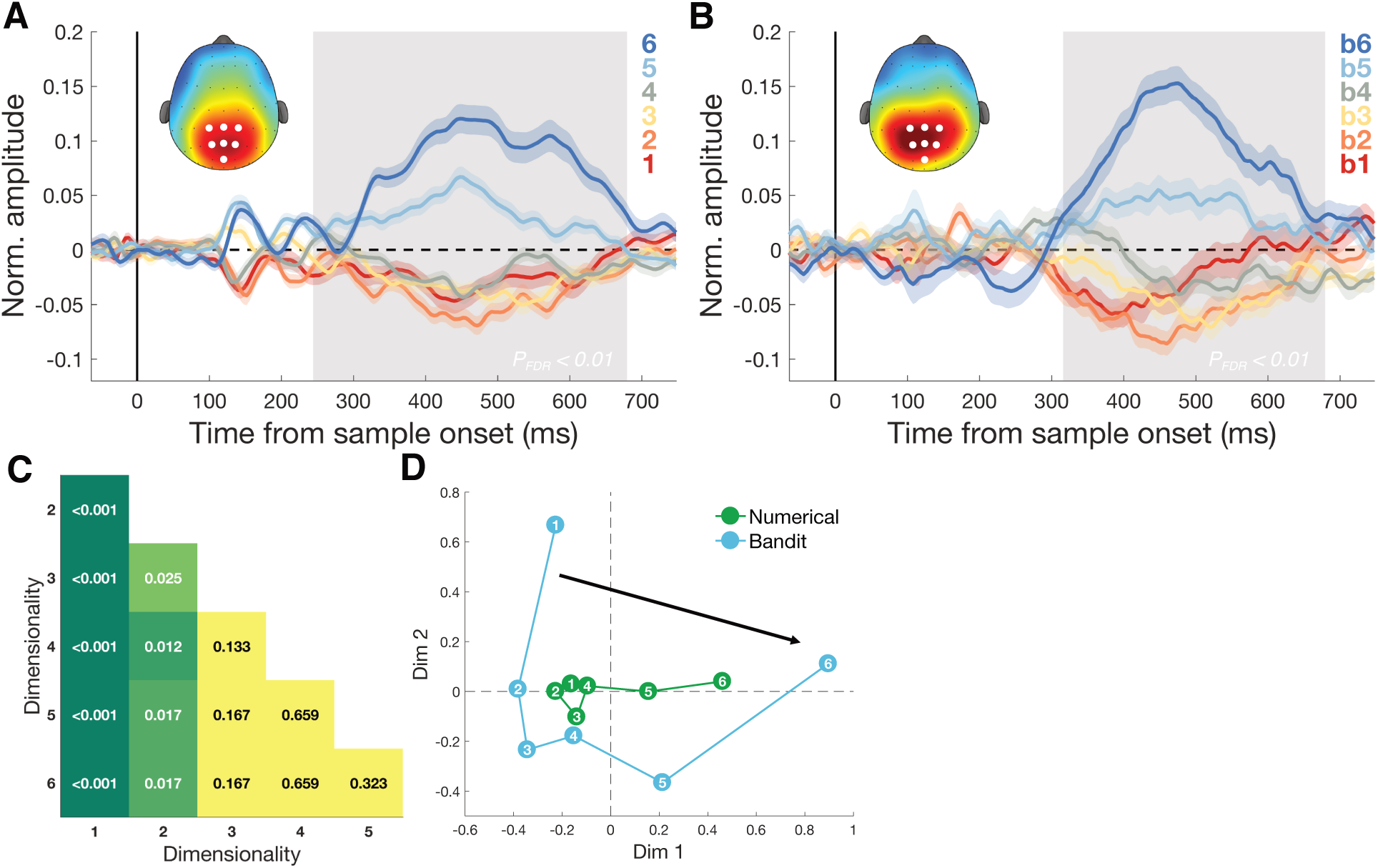
Dimensionality of magnitude representation. **(A)** Regression coefficients associated with numbers 1-6, independent of task framing (report highest/lowest) or category (blue/orange). Grey shaded area shows time of greatest disparity between signals (Kruskis-Wallis, P_FDR_ < .01). Scalp map inset shows response amplitude for digit 6 during identified time window. Colored shading represents SEM. **(B)** Equivalent analysis for bandits b1-b6 in the bandit task. Scalp map shows activation for highest subjectively valued bandit (b6). **(C)** Cross-validation after dimension reduction using SVD in each task. Each cell contains the p-value indicating the significance of the pairwise t-test comparing average cross-validation effect in the 350-600 ms time window under different dimensionalities of the data. Reduction to one dimension significantly reduced the size of the effect. **(D)** Multi-dimensional scaling (MDS) revealed two principal axes that describe the data: a magnitude axis approximately following the number/bandit order (along the black arrow) and a certainty axis distinguishing inlying (e.g. 3,4) from outlying (e.g. 1,6) numbers or bandits.

Together, these observations suggest that an abstract neural code for magnitude forms a “scaffold” for learning about the reward probabilities associated with novel stimuli. Rather than encoding stimulus value in an unstructured value function or lookup table (as is common in RL models), our data suggest that humans project available stimuli onto a one-dimensional axis that runs from “bad” to “good”. This neural axis is aligned with the mental number line, suggesting that humans recycle an abstract concept of magnitude to encode reward probabilities. Learning a structured representation of value will have the benefit of allowing new inductive inferences, such as inferring transitive preferences among economic goods without exhaustive pairwise comparison, and facilitate read-out in downstream brain areas, related to the notion of a ‘common currency’ (*30*). Although the spatial resolution of EEG is limited, our univariate data point to the parietal cortex as one locus for the low-dimensional projection of value, where neural signals for magnitude have previously been proposed to provide a conceptual bridge between different metrics such as space, time and number (*16, 17, 31, 32*).

## Acknowledgments

The authors would like to thank Zeb Kurth-Nelson for his insightful comments on an early draft of the manuscript and Mark Stokes for providing access to EEG equipment.

## Author contributions

Conceptualization: FL, CS, BS; Methodology: FL, CS, BS, HN; Software: FL, BS, CS; Validation: FL; Formal Analysis: FL; Investigation: FL; Resources: CS; Data Curation: FL; Writing – Original Draft Preparation: CS, FL; Writing – Review & Editing: FL, CS, BS, HN; Visualization: FL; Supervision: CS, BS, HN; Project Administration: CS; Funding Acquisition: CS.

## Data and materials availability

All data, code and materials to reproduce the analyses are available at https://github.com/summerfieldlab. Raw EEG data can be requested with the corresponding authors.

